# Substrate Stiffness Modulates Fibroblast Extracellular Vesicle Secretion via Mechanotransduction Pathways

**DOI:** 10.1101/2025.07.29.667440

**Authors:** Jun Yang, Lara Ece Celebi, Lauren Hawthorne, S. Gulberk Ozcebe, Pinar Zorlutuna

## Abstract

The extracellular matrix (ECM) is recognized as a key regulator of cell behavior, with its stiffness playing a crucial role in the progression of pathological conditions such as cancer and cardiovascular diseases. While extracellular vesicles (EVs) are essential mediators of intercellular communication, the influence of matrix stiffness on EV secretion remains poorly understood. This study investigates how substrate stiffness affects EV size and composition in mouse mammary and cardiac fibroblasts, the key stromal cell types in breast cancer and cardiac microenvironments. Importantly, we uncovered stiffness-tuned EV proteomic cargo, providing new insights into how mechanical cues can reprogram the signaling functions of fibroblast-derived vesicles. Our findings show that substrate stiffness significantly alters EV characteristics, with sizes increasing below stiffnesses of 20 kPa and decreasing on stiffer substrates. Mechanotransduction pathways involving p53 and thioredoxin were identified as regulators of these alterations, with thioredoxin dominating the modulation in mammary fibroblasts and p53 in cardiac fibroblasts. These results underscore the importance of ECM stiffness in modulating EV secretion and highlight candidate pathways influenced by ECM remodeling that may warrant further investigation for therapeutic relevance.

## Introduction

The extracellular matrix (ECM) is the physiologically active, non-cellular component of tissue that provides structural and biochemical support for cellular adhesion, proliferation, and intercellular communications^1^. Growing evidence has identified the ECM as a fundamental mediator in most if not all, mammalian diseases^2^. As cancer progresses, the ECM undergoes active remodeling and elicits biological cues to influence cancer cell adhesion, proliferation and migration^3^. In breast cancer development, ECM features significant alterations in composition and organization compared to the breast tissue under homeostasis^4^. Similarly, most myocardial diseases are associated with marked alterations in the ECM network^5^. The increased understanding of how ECM composition and mechanobiology modulate disease progression is important for the future development of novel therapeutics.

With growing interest in the role of ECM in various disease progression, studies highlighted matrix stiffness as a key factor in conditions such as cardiovascular diseases (CVD)^6,7^ and cancer^3,8,9^, the leading causes of death globally. Multiple studies have investigated the role of arterial stiffness in cardiovascular diseases^10^, including myocardium infarction (MI)^11^ and ischemic and nonischemic heart failure^5^. Similarly, MI^12^, obesity^13^, or age-associated^14,15^ ECM remodeling of the myocardium results in increased collagen deposition and fibrosis, leading to increase in stiffness and plays a pivotal role in the progression of cardiovascular diseases^16^, ultimately contributing to the development of heart failure by impairing myocardial compliance and contractility. On the other hand, ECM stiffness is a widely acknowledged driving force of cancer progression and tumor cell metastasis^17^. As a significant risk factor in cancer, matrix stiffness alters intracellular rheology^18^ and contributes to cancer progression^19^. In breast cancer^20,21^, the heterogeneous biophysical properties of the microenvironment alter breast cancer cell stemness^22^ and regulate breast cancer cell pro-metastatic functions^23^. Therefore, understanding these ECM-driven changes and their pathophysiological consequences is critical for developing therapeutic strategies to mitigate cardiac dysfunction and cancer progression.

As a significant and integral component of ECM^24^, extracellular vesicles (EVs) serve as mediators in multiple biochemical signaling events and transport biomolecules between cells and tissues in disease progression^25^. Many biological processes closely related to ECM remodeling and matrix stiffness, including aging and pathogenesis, have been reported to pose a significant impact on vesicle characteristics^26,27^ and biological effects^28^. For example, in a recent study, we showed that the matrix-bound vesicles (MBV) in the aged left ventricle are smaller than those bound in the young tissues^27^ and exhibit distinct expression profiles of cytokines and miRNAs, which could be a consequence of the age-related increase in ECM stiffness. However, few studies have investigated how matrix stiffness alters cellular EV secretion. A recent study revealed that ECM stiffness-tuned exosomes from breast cancer cells promote breast cancer motility through thrombospondin-1^29^. Another similar study showed that stiff EVs from cancer cells on matrices that model human breast tumors aided cancer cell dissemination and lung metastasis^30^. These studies demonstrated the differentiated effects of substrate stiffness-tuned exosomes on biological processes. However, the impact of matrix stiffness on EV secretion and the mechanisms underlying matrix stiffness-induced alterations in EV secretion remain largely unexplored. Although ECM stiffness-mediated EV alterations have not been specifically studied, existing research indicates that fibrosis-driven changes in EV content and quantity exacerbate cardiac dysfunction and contribute to heart failure progression. Tian et al. reported that cardiac fibrosis and hypertrophy drive heart failure progression by altering cardiac fibroblast-derived EV levels, leading to dysregulated signaling that exacerbates structural remodeling and dysfunction^31^. Some recent studies have focused on how substrate stiffness influences the overall quantity of EV secretion^32^ or the metastatic impact of the secreted EVs^33–35^, particularly in cancer models. In contrast, the effects of stiffness on EV size and cargo composition in stromal fibroblasts remain largely uncharacterized.

In this study, to better understand the effects of substrate stiffness on cellular EV production, we investigated the effects of substrate stiffness on mouse mammary (MMF) and cardiac fibroblast (MCF) secreted EV sizes and compositions. While the aforementioned studies have primarily focused on cancer cell-derived extracellular vesicles (EVs), matrix stiffness has been shown to significantly influence fibroblast activation within the tumor microenvironment^36^ and the myocardium^37^. This is particularly evident in the activation, senescence, and proinflammatory secretory phenotype of cardiac fibroblasts^36^. Given that mammary and cardiac fibroblasts are the predominant stromal cells in breast cancer and cardiovascular microenvironments, they represent a relevant model system for investigating how matrix stiffness modulates cellular EV secretion. Furthermore, based on the identified stiffness-tuned EV alterations, we investigated the relationship between mechanotransduction and EV secretion. To uncover potential signaling mediators, we performed proteomic profiling of EV cargo and mapped differentially expressed proteins to known mechanotransduction pathways. With pathway analysis and protein inhibition assays, we successfully identified signaling pathways and corresponding effector proteins that exerted a significant impact on how substrate stiffness affected fibroblast EV secretion.

## Materials and Methods

### Polyacrylamide Gel Preparation and Characterization

For PA gel preparation, an estalished protocol was adapted^38^. Briefly, acrylamide solution (40% w/v, Thermo Fisher, CA), N,N’-methylenebisacrylamide (1% w/v bis-AA, VWR, OH), HEPES buffer (10mM, Gibco, NY) and ammonium persulfate (10% w/v APS, VWR, OH) stock solutions were prepared in distilled water. Acrylamide and bis-AA solutions were mixed in desired ratios (Table 1) in HEPES buffer. Polymerization was initiated with the addition of 1/100 total volume of APS and 1/1000 total volume of tetramethylethylenediamine (TEMED, Sigma) into the Acrylamide/bis-AA solution. Then, the solution was loaded into the casting frame (BioRad) and gelation took place for 30 minutes. After the gelation, each gel was transferred in PBS and kept at 4°C until use. Mechanical properties of the PA gels were measured with Optics11Life Chiaro nanoindenter with probes of 0.025 N/m stiffness and 15-25 μm tip radius. Ten spatially distinct nanoindentation measurements were conducted per gel (technical replicates), across three independently fabricated gels per stiffness (biological replicates), totaling 30 measurements per condition.

**Table 1.** Polyacrylamide Gel Composition.

|  |  |  |  |  |  |
| --- | --- | --- | --- | --- | --- |
| Acrylamide ratio | 7.5% | 7.5% | 7.5% | 7.5% | 10% |
| Bis-AA ratio | 0.04% | 0.08% | 0.16% | 0.32% | 0.32% |

### Polyacrylamide Gel Coating

PA gels were molded into cylinders using a biopsy punch. Then, the surface of PA gels was covered with sulfosuccinimidyl-6-(4’-azido-2’-nitrophenylamino)-hexanoate (sulfo-SANPAH 50mM in 10 mM HEPES, pH 8.0, Covachem, IL) and placed under UV (90 mW/cm2) for 120 seconds. After that, constructs were washed with PBS and the functionalization was repeated. Lastly, gels were triple-washed and transferred into the 96-well plate and coated with 0.2 mg/mL type I collagen in 0.1% acetic acid overnight.

### Cell Culture and Seeding

Mouse cardiac fibroblast cell line purchased from iXCells Biotechnologies (Cat#10MU-015, San Diego, CA, USA) was cultured in cell growth medium (DMEM [high glucose] medium supplemented with 10% FBS [Thermo Fisher Scientific] and 1% penicillin/streptomycin [Corning]) with SD208. Mouse mammary fibroblast cell line purchased from Cell Biologics (Cat# NC1966214, Chicago, IL, USA) was cultured in a commercial fibroblast medium kit (Cell Biologics, Cat#M2267, Chicago, IL, USA) with which includes basal medium, fetal bovine serum, fibroblast growth factor, hydrocortisone, and antibiotic-antimycotic solution. Cells were maintained in culture until 90% confluency in a CO_2_ incubator at 37 °C and 95% humidity. They were passaged using 0.25% trypsin-EDTA, reconstituted in cell growth media, and seeded in culture flasks or plates.

### Phalloidin Staining

Phalloidin staining was performed on cells cultured on coated PA gels for 2 days after observed cell attachment. Fibroblasts were fixed with 4% Paraformaldehyde (PFA) and stained with Alexa Fluor 594 tagged anti-phalloidin antibody and 1 μg/mL DAPI. Stained samples were visualized microscopically with an inverted microscope.

### α-SMA and Vimentin Staining

α-smooth muscle actin (α-SMA) and Vimentin staining was performed on fibroblasts cultured on coated PA gels for 3 days, after initial cell attachment. The cells were fixed with 4% paraformaldehyde (PFA), permeabilized with 0.1% Triton-X, and blocked using 4% goat serum. Subsequently, the cells were incubated with primary antibodies against α-SMA (Abcam, ab7817) and Vimentin (Abcam, ab92547) in 4% goat serum overnight. The following day, the PA gels were washed with PBS and incubated with secondary antibodies, Goat Anti-Mouse IgG H&L Alexa Fluor 647 (Abcam, ab150115) and Alexa Fluor 488 (Abcam, ab150077), in 4% goat serum for 3 hours at 4°C. After incubation, constructs were washed again and stained with 1 μg/mL DAPI. PA gels were then carefully taken out from 96-well plates, placed on thin glass slides, and visualized using Airyscan and fluorescence microscopy. Alpha-SMA-expressing cells were quantified using ImageJ, normalized to the cell count (DAPI), and graphed utilizing GraphPad Prism.

### Vesicle Isolation

Cell medium was collected every two days after observed cell adhesion. Each sample contains cell medium from 6 wells in a 96 well plate. The collected cell medium was subjected to successive centrifugations at 500 x g for 10 min, 2,500 x g for 20 min, and 10,000 x g for 30 min. Each centrifugation step was performed three times to ensure the removal of collagen fibril remnants. The fiber-free supernatant was then centrifuged at 100,000 x g with tabletop ultracentrifuge (Beckman Coulter) for 70 min at 4 °C. The pellets were stored at −80 °C or resuspended in PBS for immediate use.

### Transmission Electron Microscopy Imaging

Isolated vesicle morphology characterization was confirmed with TEM imaging. Briefly, vesicle samples (n=3) isolated with ultracentrifugation were resuspended and incubated in 2.5% glutaraldehyde for 1 hour at room temperature and loaded on plasma plasma-cleaned carbon-coated copper grid (Polysciences). Fixed samples were incubated on the grids for 20 min at room temperature. The samples were washed with deionized water and vacuumed for 10 min to dry. MBV samples were incubated in vanadium solution (Abcam) for 15 min at room temperature for negative staining. The samples were then washed with deionized water and vacuum-dried. TEM imaging was performed at 120 kV.

### Vesicle Size Measurement

EV samples were collected through ultracentrifugation methods and washed with PBS. Pellets were resuspended in 120 μL deionized water and dynamic light scattering measurements were performed with an Omni Nanobrook.

Nanoparticle tracking analysis (NTA) was performed with a NanoSight NS3000 instrument (Malvern). Size distribution and particle concentration of MBVs were determined by Brownian motion measurement (n=3).

### Surface Marker Quantification

Quantification of surface markers of vesicle samples was performed with ExoELISA-ULTRA Complete Kit for CD9 and CD63 detection (SBI). Isolated MBVs were resuspended in PBS and added into protein-binding 96-well plates at 5 μg per well. Plates were incubated at 37 °C for an hour for protein binding and washed 3 times with 1x wash buffer. Plates were then treated with corresponding primary and horseradish peroxidase-labelled secondary antibodies and super-sensitive tetramethylbenzidine substrates according to manufacturer’s instructions. Reactions were terminated with stop buffer and quantitative results were measured at 450 nm with a plate reader (Wallac 1420).

### Mechanosensing Intervention

Blebbistatin (1-phenyl-1,2,3,4-tetrahydro-4-hydroxpyrrolo[2,3-b]-7-methylquinolin-4-one) was dissolved in DMSO at a concentration of 10 mM. Rock-Inhibitor (Y-27632, (+)-(R)-trans-4-(1-aminoethyl)-N-(4-pyridyl)cyclohexanecarboxamide dihydrochlorate) was dissolved in DMSO at a concentration of 10 mM. Mechanosensing pathways were blocked through the addition of 50 μM myosin inhibitor blebbistatin or 10 μM ROCK inhibitor Y-27632.

### Proteomics

EV samples were collected through ultracentrifugation methods and washed with PBS. Pellets were resuspended in 30 μL PBS and protein concentration was assessed with BCA. 30 μg protein of each sample was transferred to a fresh tube, reduced in SDS buffer with 20 mM dithiothreitol, and heated at 65 C for 20 min. Cysteines were alkylated by adding 40 mM iodoacetamide and incubated in the dark for 30 min. The processed samples were then added into the S-Traps and incubated in trypsin at 37 C overnight. The proteins were eluted in acetonitrile, dried, reconstituted, and desalted with C18 ziptips (Thermo Fisher). Bottom-up proteomics was performed with a Thermo Q-Exactive HF with 1 μL injection.

Ingenuity pathway analysis (IPA) was carried out with Metacore online platform.

### Western Blot

Mouse mammary and cardiac fibroblasts were cultured on 10, 20, 60 kPa substrate for five days and lysed using RIPA buffer (50 mM Tris-HCl, pH 7.4, 150 mM NaCl, 1% NP-40, 0.5% sodium deoxycholate, 0.1% SDS) supplemented with protease and inhibitors (Abcam). Protein concentration was determined using the BCA protein assay kit (Thermo Fisher Scientific), following the manufacturer’s instructions. Equal amounts of protein (20 μg) were denatured and loaded for western blot. Samples were probed against thioredoxin (TRX) (Abcam ab273877, 1:1000), talin-1 (Abcam ab108480, 1:1000), JunB (Abcam ab128878, 1:200), p53 (Abcam ab26, 1:1000), c-Myc (Abcam ab32072, 1:1000), beta-catenin (Abcam ab196204, 1:5000) and internal control with beta-actin (Abcam ab8227, 1:1000). Where necessary, membranes were stripped using stripping buffer (Thermo Fisher Scientific) for 30 minutes at room temperature, re-blocked, and re-probed with different primary antibodies.

### Pathway Inhibition Study

Protein inhibitors utilized in the functional assays were Pifithrin-α (PFTα) HBr (Pifithrin-α hydrobromide dissolved in DMSO, Selleckchem S2929), MYCi975 (NUCC-0200975 dissolved in DMSO, Selleckchem S8906) and PX-12 (1-methyl propyl 2-imidazolyl disulfide dissolved in DMSO, Selleckchem S7947). MMFs and cardiac fibroblasts seeded on five different stiffness substrates were treated with the aforementioned protein inhibitors 24 hours after seeding. After 48 hours of inhibitor treatments (24 hours treatment for PX-12), cells were switched to exosome-free medium, conditioned medium was collected every 48 hours, and EVs were extracted through ultracentrifugation methods and washed with PBS for DLS measurements.

### Statistical Analysis

Data were analyzed for statistical significance with Prism 10 (Graphpad). Two-tailed unpaired student’s t-test was applied to compare the difference between two groups and one-way ANOVA followed by Tukey’s HSD correction was performed to compare the differences between multiple groups. Outliers were identified using the ROUT method with Q = 1% and eliminated. Data are presented as the mean ± standard deviation (SD).

## Results and Discussion

### EV sizes alter with substrate stiffness

To comprehensively understand the impact of matrix stiffness on fibroblast EV secretion, a wide series of physiologically and pathologically relevant substrate stiffnesses was selected. Mechanically tunable polyacrylamide (PA) hydrogels are widely utilized as cell culture substrates to investigate how cells perceive and adapt to the physical properties of their microenvironment^39,40^. Young’s modulus of polyacrylamide (PA) gels with different tris-bis compositions was assessed using nanoindentation, which was chosen for its ability to measure mechanical properties at the cellular length scale (10–20 µm), directly reflecting how cells sense substrate stiffness. Substrate stiffness in this study ranged from around 1 kPa, the stiffness of healthy young decellularized breast tissue scaffolds, to over 60 kPa, the stiffness value of left ventricular tissues with pathological conditions. The selected stiffness series also includes the stiffness of the solid breast tumor (> 25 kPa)^23^, as well as healthy myocardium tissues (10 kPa), and border and scar regions (∼60 kPa) of post-MI cardiac tissues^41^.

Cell adhesion of both mouse mammary and cardiac fibroblasts was confirmed with phalloidin staining (Figure 1C), demonstrating the cell morphology when cultured on the collagen-coated PA gels. To assess differences in fibroblast mechanosensitivity, we performed immunofluorescence staining for α-SMA, a myofibroblast marker indicative of activation (Supplementary Figure 1). As expected from literature, α-SMA expression was shown to increase with stiffness^42^, particularly in MMFs, where substrate over 60 kPa induces a significant upregulation of α-SMA compared to lower stiffness conditions. The increased α-SMA expression suggests increased myofibroblast activity, ECM remodeling, and a potentially more fibrotic microenvironment. In contrast, MCFs exhibit relatively stable α-SMA levels across conditions, suggesting a differential mechanosensitive response between these fibroblast populations. The supernatant was collected from MMFs and MCFs cultured on substrates with varying stiffness, and EVs were isolated through ultracentrifugation as reported previously^26,27,43^.

**FIGURE 1.**
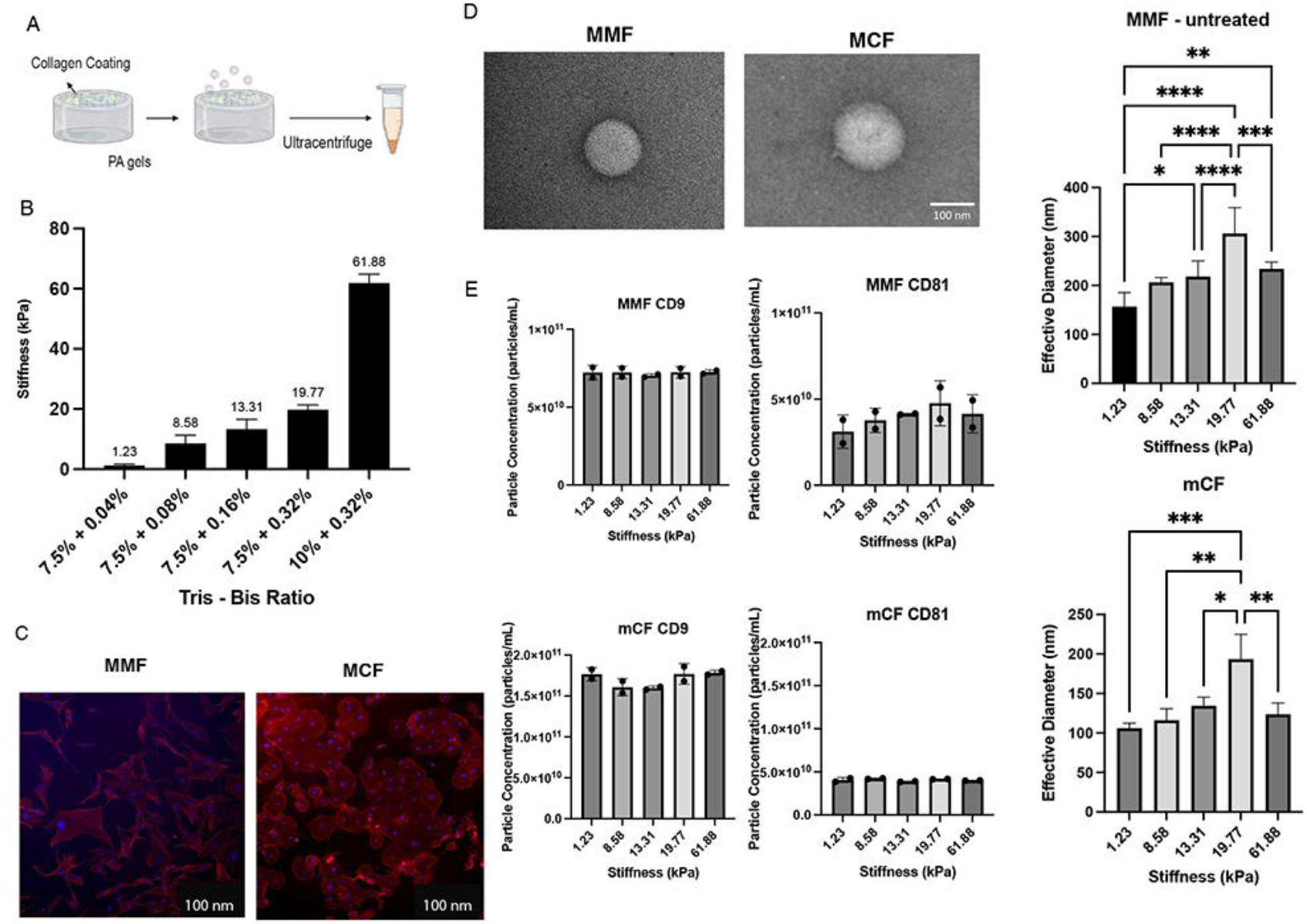
Substrate stiffness alters the sizes of fibroblast-secreted EVs. A) Schematics of EV generation methods. B) Stiffness of PA gels with selected composition. C) Phalloidin staining confirmed cell attachment of MMF (left) and MCF (right) to collagen-coated PA gels. D) Representative TEM images confirmed morphology of MMF and MCF secreted EVs. E) Size and surface marker expression of EVs from MMFs and MCFs cultured on different substrate stiffness (n=6). Data presented as the mean ± SD. ANOVA followed by Tukey’s post hoc was applied for statistical significance. *p<0.05, **p<0.01, ***p<0.001, and ****p<0.0001.

TEM imaging with a negative stain (Figure 1D) confirmed the morphology of the isolated EVs to be spherical vesicles over 100 nm in diameter. DLS size measurements demonstrated that sizes of the EVs secreted by both MMFs and MCFs increased with stiffness up to 20 kPa, where they reached a significantly larger size compared to other conditions. MMF-secreted EVs presented an average diameter of 309.1 ± 52.5 nm from MMFs cultured on 20 kPa substrate and average diameters between 150-200 nm from MMFs cultured on other stiffness conditions (Figure 1E). Similarly, MCF secreted EVs had an average diameter of 191.2 ± 35.4 nm from MCFs cultured on 20 kPa PA gels while they had average diameters between 100-140 nm from MCFs cultured on PA gels of other stiffness. These observed EV size changes, particularly the reduction in diameter under high-stiffness conditions, may be related to the aforementioned stiffness-induced phenotypic alterations in fibroblasts. However, further investigation is needed to clarify the link between fibroblast activation and EV secretion dynamics. One possible explanation for this biphasic EV size trend is the differential activation thresholds of key mechanotransducers. A substrate stiffness of 20 kPa represents a physiologically relevant transitional zone (border zone) between healthy (10 kPa) and fibrotic myocardium (∼60 kPa), and mirrors early matrix remodeling during tumor progression^41,44^. For instance, moderate stiffness may optimize cytoskeletal tension and intracellular vesicle trafficking via ROCK1 and TRX activation, promoting larger vesicle formation. At higher stiffness levels, excessive mechanical stress may activate alternative signaling cascades such as actin reorganization and YAP/TAZ-mediated transcription. These complex mechanotransduction responses may underlie the non-linear regulation of EV size in response to matrix stiffness.

Surface marker quantification with ELISA further confirmed similar expression levels of EV surface markers CD9 and CD81 on EVs secreted by cells cultured on different substrate stiffness. Interestingly, the EV concentration per sample measured by nanoparticle tracking analysis (NTA) from all tested conditions showed no significant difference compared to the EV concentration measured by CD9 ELISA (Supplementary Figure S2), indicating that most of the secreted EVs are CD9 positive. This demonstrates that substrate stiffness does not affect the amount of EV secreted, and that no significant EV fusion or fission events occurred under these conditions. On the other hand, in both types of fibroblasts secreted EVs, CD81 expression was lower than CD9 expression level, while CD81 expression was also not affected by substrate stiffness. Characterization of the isolated vesicles demonstrated that increased substrate stiffness alters vesicles towards larger sizes on softer substrates (< 20 kPa) but smaller sizes on stiffer substrates (> 20 kPa). This finding suggests that substrate stiffness plays a role in modulating fibroblast-secreted vesicle characteristics. However, different substrate stiffnesses do not alter secreted vesicle concentration or surface marker expression levels, similar to the consistent EV surface marker expression observed in healthy aged and young tissues in previous studies^26,27^.

### Mechanotransduction inhibition attenuates the effect of substrate stiffness on EV secretion

To test the observed changes in vesicle sizes were influenced by stiffness, experiments were conducted to inhibit mechanotransduction pathways. MMFs and MCFs were treated with two well-characterized inhibitors that target key regulators of the cellular contractility and mechanotransduction pathways, blebbistatin, a myosin II inhibitor, and Y-27632, a highly potent and selective ROCK inhibitor. Following treatment, EVs secreted by both cell types did not exhibit size changes in response to substrate stiffness (Figure 2A-2B). With blebbistatin treatment inhibiting myosin II, a well-studied bipolar actin-based motor protein, MMFs cultured on any substrate stiffness secreted EVs with an average diameter of around 150 nm, while all MCFs secreted EVs around 120 nm. On the other hand, with Y-27632 treatment blocking ROCK I and ROCK II, kinases that mainly regulate cell shape and movement, both MMFs and MCFs cultured on the substrate with varied stiffness secreted EVs with an average diameter of around 200 nm. This further indicated that the changes in EV sizes observed in Figure 1E are indeed due to the varied substrate stiffness. Furthermore, the different sizes observed in blebbstatin-treated fibroblast-secreted EVs and Y-27632-treated fibroblast-secreted EVs indicate that inhibition of different mechanosensing pathways may affect EV secretion differently, suggesting that substrate stiffness influences EV secretion via multiple, pathway-specific mechanisms. While both treatments eliminated the stiffness-dependent differences in EV size, blebbistatin consistently reduced EV size whereas Y-27632 increased it, suggesting that myosin II and ROCK inhibition may affect membrane curvature, actin dynamics, or vesicle scission in divergent ways that differentially regulate vesicle formation and release^45,46^.

**FIGURE 2.**
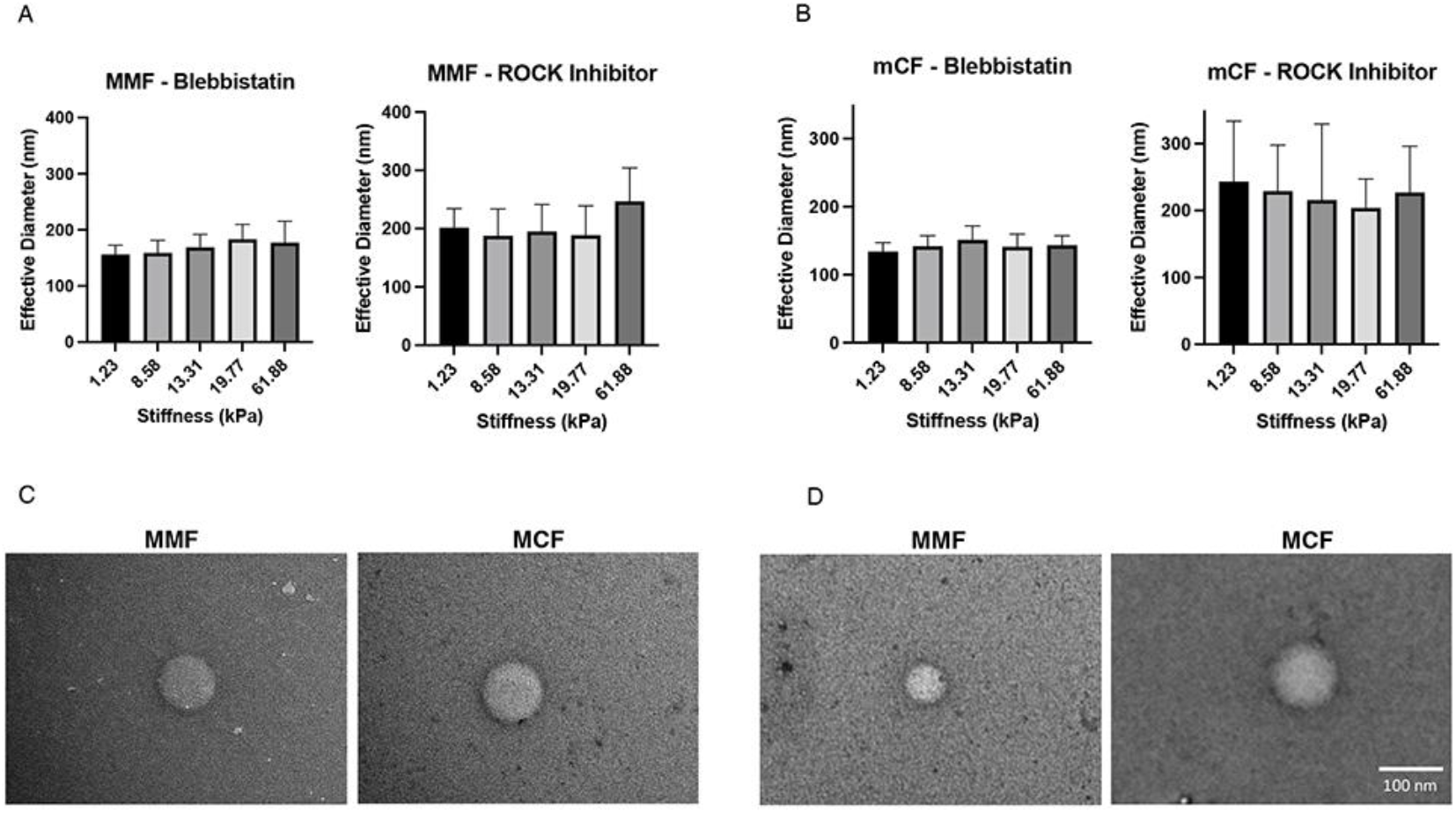
Mechanotransduction inhibition attenuates the effect of substrate stiffness on fibroblast-secreted EV sizes. A) inhibition of mechanotransduction inhibited stiffness-induced EV size changes in MMFs (n=6). B) Inhibition of mechanotransduction inhibited stiffness-induced EV size changes in MCFs (n=6). C) TEM images confirmed morphology of MMF-EVs with mechanotransduction inhibition. D) TEM images confirmed morphology of MCF-EVs with mechanotransduction inhibition. Data presented as the mean ± SD. ANOVA followed by Tukey’s post hoc was applied for statistical significance.

Meanwhile, TEM imaging confirmed that MMF and MCF-secreted EVs maintained their spherical morphology (Figure 2C-2D), indicating that while mechanotransduction inhibition attenuates the effects of substrate stiffness on EV sizes, it does not alter vesicle morphology. Notably, although mechanosensing was inhibited, fibroblasts can still secrete EVs with perserved morphology, indicating that their basal EV production and potential for intercellular communication remain intact.

### Substrate stiffness alters the proteome of fibroblast-secreted EVs

Cells utilize EVs to convey biomolecules such as proteins and miRNAs to communicate with the microenvironment and other cells. While substrate stiffness alters the size of fibroblast-secreted EVs, label-free proteomics quantification analysis of these EVs showed that substrate stiffness also affects the protein contents carried by these fibroblast-secreted EVs (Supplementary Figure S3). Among the proteins identified in the EV samples, some exhibited expression changes in response to substrate stiffness, while others showed trends that paralleled the observed alterations in EV size (Figure 3A). Notably, several redox-associated proteins, including thioredoxin, thioredoxin-dependent peroxidase reductase, and peroxiredoxin-5, were enriched in MMF-derived EVs, with their expression levels particularly changing with substrate stiffness. These findings suggest that redox signaling components are selectively packaged into EVs under specific mechanical cues.

**FIGURE 3.**
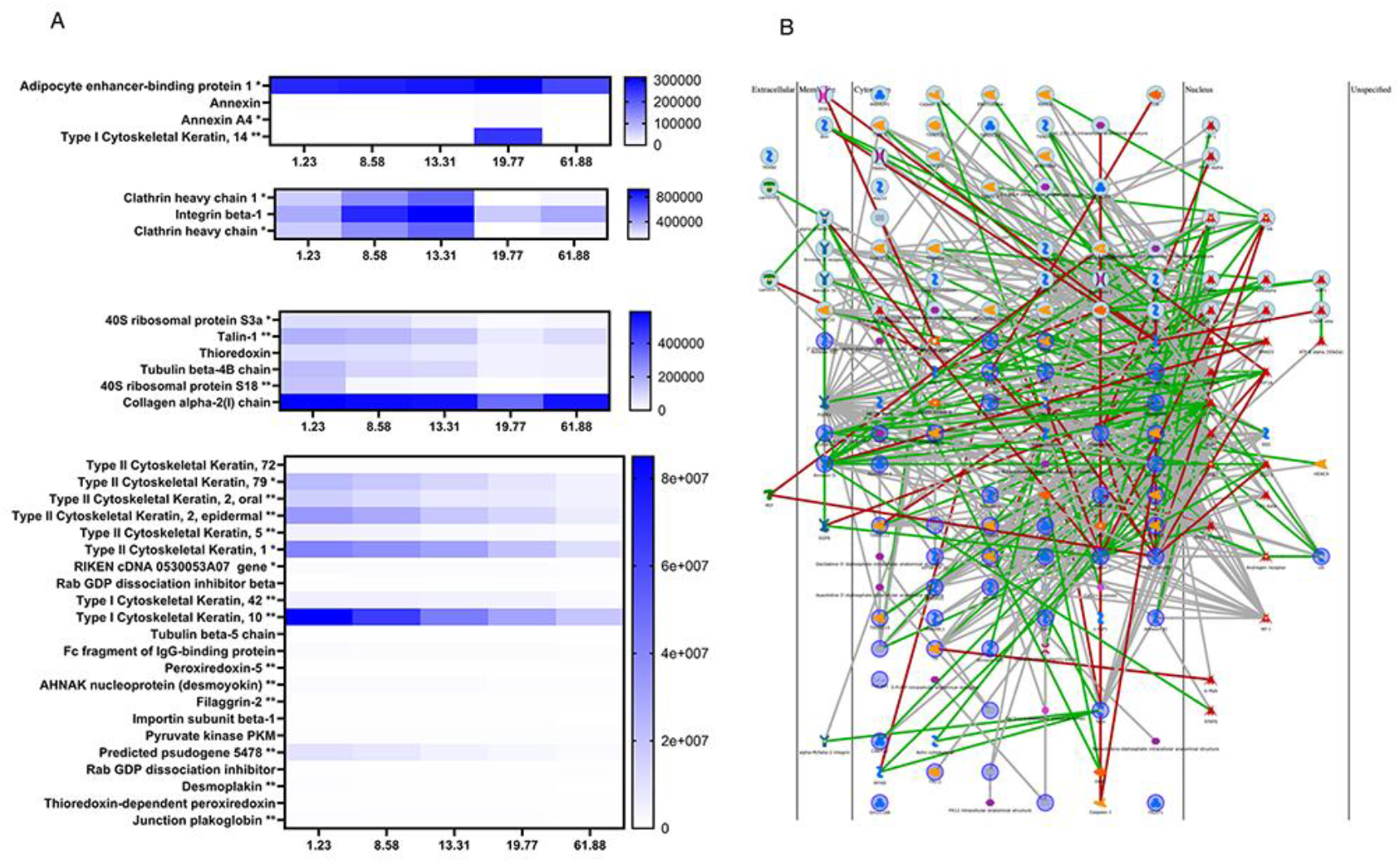
Substrate stiffness alters the proteome of EVs secreted by MMF and MCF. A) Proteomic analysis identified specific proteins within fibroblast-secreted EVs with expression levels altered by substrate stiffness (n=3). MMF-specific proteins are labeld with * and MCF-specific proteins are labeled with ** in the heatmap. B) Pathway analysis of the identified proteins highlights key biological processes affected by these changes.

IPA pathway analysis of these proteins identified multiple signaling pathways potentially linking the observed alterations in protein expression (Figure 3B). To further elucidate these connections, we cross-referencing our findings with existing literature on the effects of substrate stiffness on the cellular proteome and proteins associated with EV biogenesis. This comparative analysis, combined with further pathway analysis revealed that several of these differentially expressed proteins may play critical roles in mechanotransduction processes. Furthermore, since the identified proteins are involved in both ESCRT-dependent and ESCRT-independent EV secretion pathways, their differential expression with stiffness suggests that substrate stiffness may regulate EV secretion via multiple, mechanosensitive mechanisms. These findings raise important questions about how stiffness-mediated changes in EV size and cargo influence downstream cellular behaviors. Future studies will be needed to assess how EVs secreted from fibroblasts cultured on matrices of varying stiffness affect recipient cell proliferation, migration, activation, and gene expression profiles, which may shed light on how mechanical cues regulate intercellular signaling via vesicles.

Among the identified proteins that are involved in mechanotransduction pathways, TRX(Figure 4A), Talin (Supplementary Figure S4A), JunB (Supplementary Figure S4B) are closely related to the ESCRT-dependent EV biogenesis pathways, while p53 (Figure 4B), c-Myc (Figure 4C) and beta-catenin (Supplementary Figure S4C) are involved in both ESCRT dependent pathways via RAS and ESCRT independent pathways via sphingomyelinase.

**FIGURE 4.**
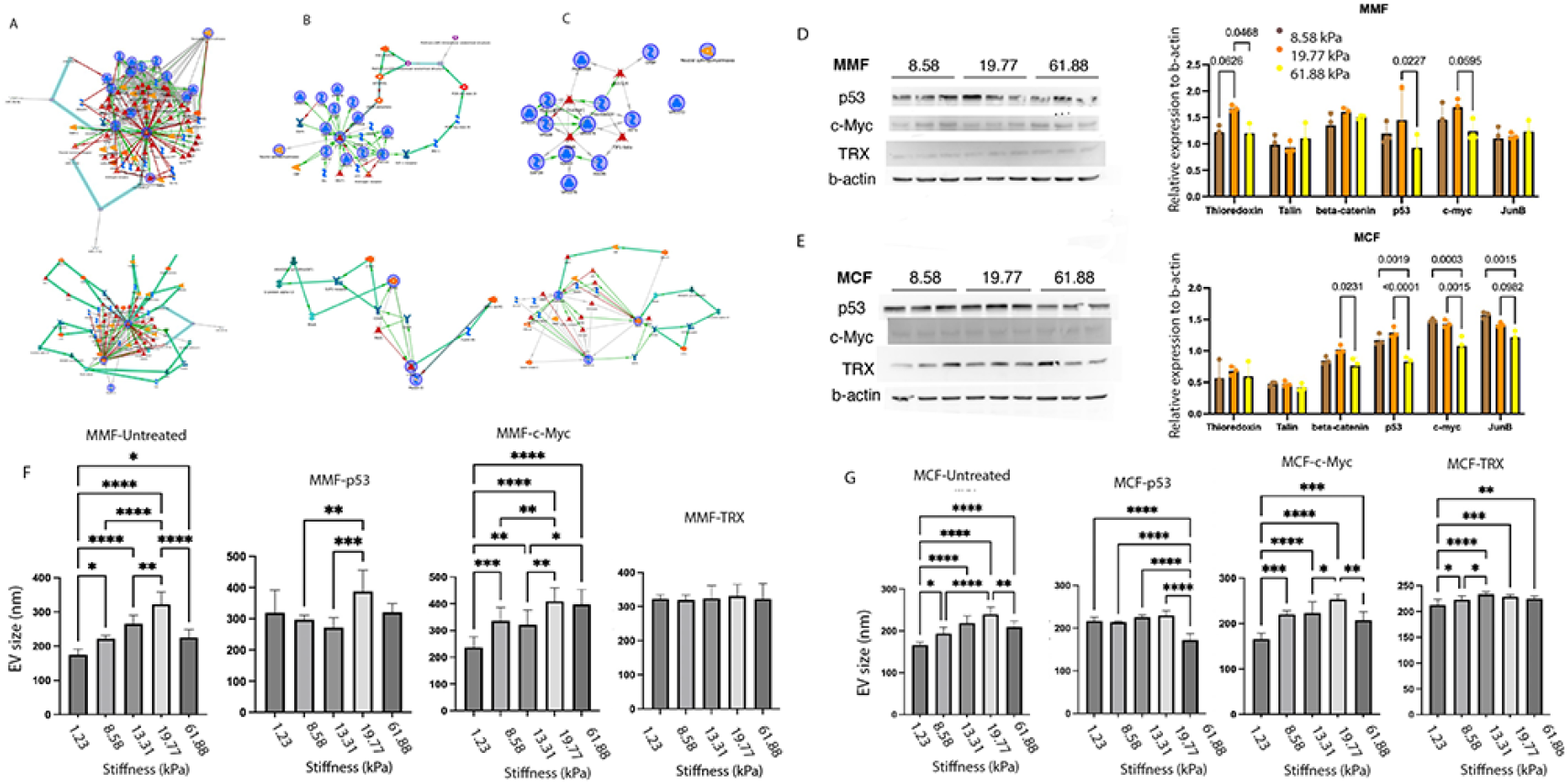
Mechanism studies identified pathways linking cellular mechanotransduction and EV secretion. A) Pathway analysis of TRX with EV biogenesis and mechanotransduction pathways. B) Pathway analysis of p53 with EV biogenesis and mechanotransduction pathways. C) Pathway analysis of c-Myc with EV biogenesis and mechanotransduction pathways. D) Western blot of identified protein expression in MMFs. E) Western blot of identified protein expression in MCFs. F) Stiffness-induced EV size differences with identified protein inhibited (n=3). G) Stiffness-dependent EV size changes following inhibition of identified mechanotransduction proteins in MCFs (n=3). Data presented as the mean ± SD. ANOVA followed by Tukey’s post hoc was applied for statistical significance. *p<0.05,**p<0.01, ***p<0.001, and ****p<0.0001.

### Substrate stiffness-altered proteins are related to EV secretion pathways

To investigate how substrate stiffness affects mechanotransduction-related protein expression, we performed western blot analysis on fibroblast cell lysates targeting proteins previously identified in our proteomic and pathway analysis as key regulators of EV biogenesis and stiffness-responsive signaling. Among the assessed proteins (Figure 4D-E), p53 and c-Myc expression levels were significantly altered with stiffness in both mouse mammary and cardiac fibroblasts. Both p53 and c-Myc showed elevated expression in mammary fibroblasts cultured on 20 kPa substrate, the same culture condition that increased EV size. In cardiac fibroblasts, p53 also showed elevated expression in cells cultured on soft substrate and lower expression on 60 kPa substrate, while c-Myc demonstrated lowered expression in cells cultured on stiffer substrates. Meanwhile, TRX expression levels were also elevated significantly in mammary fibroblasts cultured on 20 kPa substrate compared to softer (p=0.0626) and stiffer (p=0.0468) substrates.

To evaluate whether substrate stiffness poses an impact on EV secretion through these identified targets, protein inhibition was performed on fibroblasts cultured on substrates with different stiffness to examine the change in EV production and properties. Pifithrin-α (PFTα) HBr has been proven to effectively inhibit p53-dependent transactivation of p53-responsive genes in multiple cell lines with 10 μM treatment for 48 hours^47^. MYCi975 is an established Myc protein inhibitor with proven efficacy with 10 μM treatment for 48 hours^48,49^. MYCi975 selectively inhibits MYC through disruption of MYC/MAX interaction, promoting MYC T58 phosphorylation and MYC degradation, and impairs MYC-driven gene expression^50^. PX-12 is a potent thioredoxin-1 (Trx-1) inhibitor by irreversible thioalkylation of Cys73 of Trx-1, which has been shown to present significant TRX inhibition with 5 μM treatment for 24 hours^51^.

DLS results indicated that TRX inhibition demonstrated similar effects on MMF EV secretion as mechanotransduction inhibition. With 24-hour PX-12 treatment, MMFs cultured on substrates with different stiffness secreted EVs with no significant differences in size (Figure 4F). This indicates that in MMFs, EV secretion is mediated by mechanotransduction through the c-Myc/TRX axis, operating within the c-Myc/c-Jun signaling pathway via ROCK1. Blocking of the upstream protein c-Myc weakened the effects of substrate stiffness on EV secretion but did not overwrite the effects completely, indicating the effector protein was TRX in the signaling pathway.

In MCFs (Figure 4G), TRX inhibition only showed blocking of the stiffness-tuned size alterations on substrates over 13 kPa. On the other hand, p53 inhibition blocked the stiffness-induced size alterations on softer substrates below 20 kPa. Instead of being fully modulated by the ROCK1-regulated c-Myc/TRX axis, stiffness-tuned MCF EV secretion may also be controlled by myosin IIA mediated p53 functions and ROCK1-modulated p53 signaling pathway on softer substrates.

Stiffness-mediated EV size alterations in both types of fibroblasts are regulated by ROCK1-regulated TRX signaling. A few recent studies investigated the interaction between the Hippo-YAP/TAZ signaling pathway with redox signaling^52^ and reactive oxygen species accumulation^53^. The differential involvement of TRX and p53 in MMFs and MCFs suggests a redox-sensitive regulatory mechanism influenced by substrate stiffness, which may contribute to tissue-specific fibroblast behavior in fibrosis and tumor progression^54^.Our results further confirmed the significance of redox signaling in mechanotransduction and its impact on fibroblast cellular processes, including EV secretion, which will further convey fibroblast secretomes to affect further biological processes.

Meanwhile, recent studies have shed light on the mechanism of p53 regulation of cardiac fibroblast proliferation and activation^55^, but no similar involvement has been identified in most other types of fibroblasts, including mammary fibroblasts. Recruitment of p53 by promyelocytic leukemia protein in response to TGFβ1-induced myocardial fibrosis has been reported to be a new mechanism of cardiac fibroblast activation^56^. The controlling role of p53 in cardiac fibroblast accumulation and fibrotic responses^54^ could explain the unique influences of p53 on cardiac fibroblast EV secretion. Emerging evidence indicates that EV biogenesis is closely linked to lysosomal trafficking and autophagy, both of which are responsive to cytoskeletal tension and redox state. Given the known roles of TRX and MYC in regulating oxidative and metabolic stress, our findings suggest that stiffness-tuned EV secretion may also involve mechanosensitive modulation of vesicle trafficking via lysosomal or autophagic routes^57^. This study identified the ROCK1-mediated c-Myc/TRX axis and mechanotransduction-related p53 signaling as important effectors in stiffness-tuned EV secretion changes. Further investigations are needed to better understand the confirmed protein targets examined in this study.

## Conclusion

In this study, we shed light on how substrate stiffness influences the EV secretion from fibroblasts, with observed effects in both mammary and cardiac fibroblasts. We demonstrated that changes in substrate stiffness altered EV sizes and protein contents, with EV sizes increasing with stiffness below 20 kPa but decreasing with stiffer substrates. Pathway analysis with cellular and EV protein expression showed that the observed EV size alterations are modulated through mechanotransduction pathways, particularly those involving ROCK1, p53, and TRX. Our results indicate that while stiffness-mediated changes in EV size are consistent across both fibroblast types, the underlying mechanisms diverge in terms of the specific proteins involved, with TRX regulating EV secretion in mammary fibroblasts, and p53 playing a more prominent role in cardiac fibroblasts. These findings highlight the critical role of matrix stiffness in regulating fibroblast function and suggest that mechanotransduction pathways may represent novel therapeutic avenues for diseases involving ECM remodeling, such as cancer and cardiovascular diseases. While our findings support associations between stiffness and EV secretion, additional mechanistic validation is needed before therapeutic implications can be confirmed. Future research will be essential to further elucidate the full scope of the molecular mechanisms driving stiffness-induced EV secretion and its implications in disease progression

## Supporting Information

**Supplementary Figure S1.**
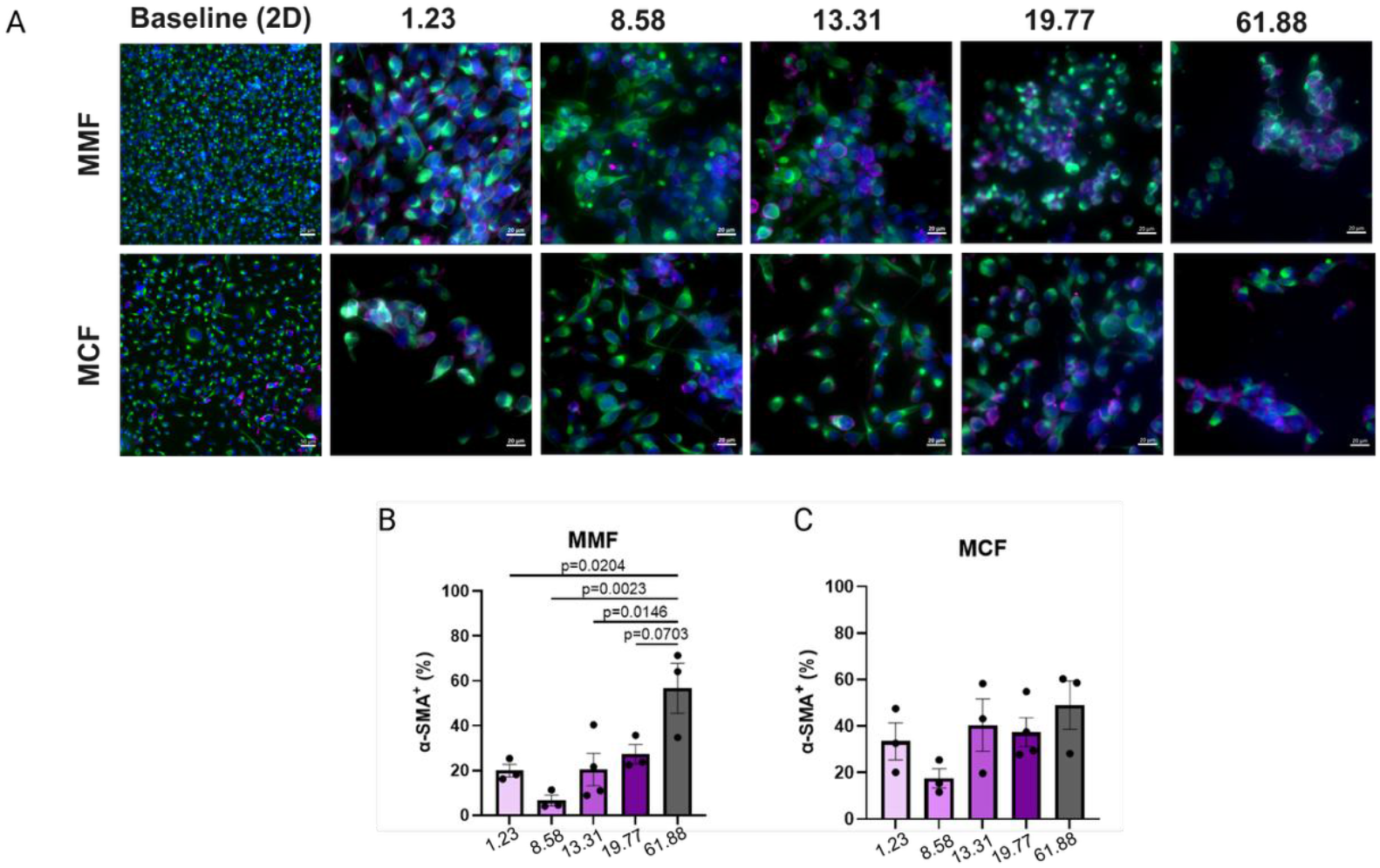
MMF and MCF activation assessment. A) α-SMA and vimentin staining of MMFs and MCFs cultured on substrates with different stiffness, baseline indicating the MMF and MCF seeded in a monolayer.B)Quantified MMF α-SMA expression levels. C) Quantified MCF α-SMA expression levels. Data presented as the mean ± SD. ANOVA followed by Tukey’s post hoc was applied for statistical significance.

**Supplementary Figure S2.**
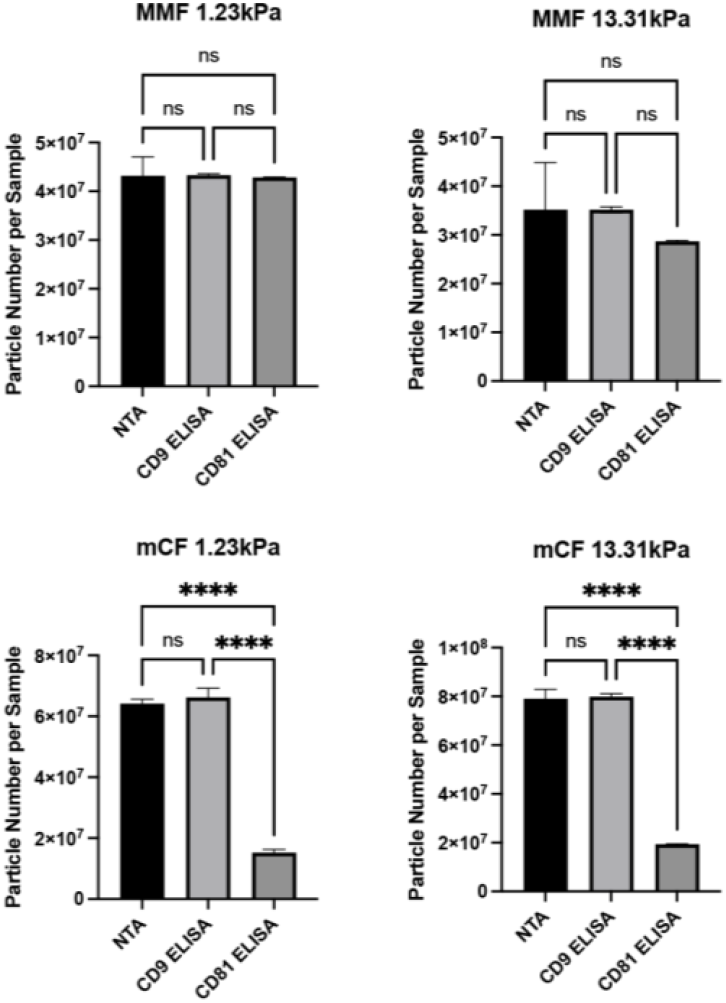
Comparison between NTA-measured EV concentration and surface marker ELISA-measured EV concentration. Data presented as the mean ± SD. ANOVA followed by Tukey’s post hoc was applied for statistical significance. *p<0.05, **p<0.01, ***p<0.001, and ****p<0.0001.

**Supplementary Figure S3.**
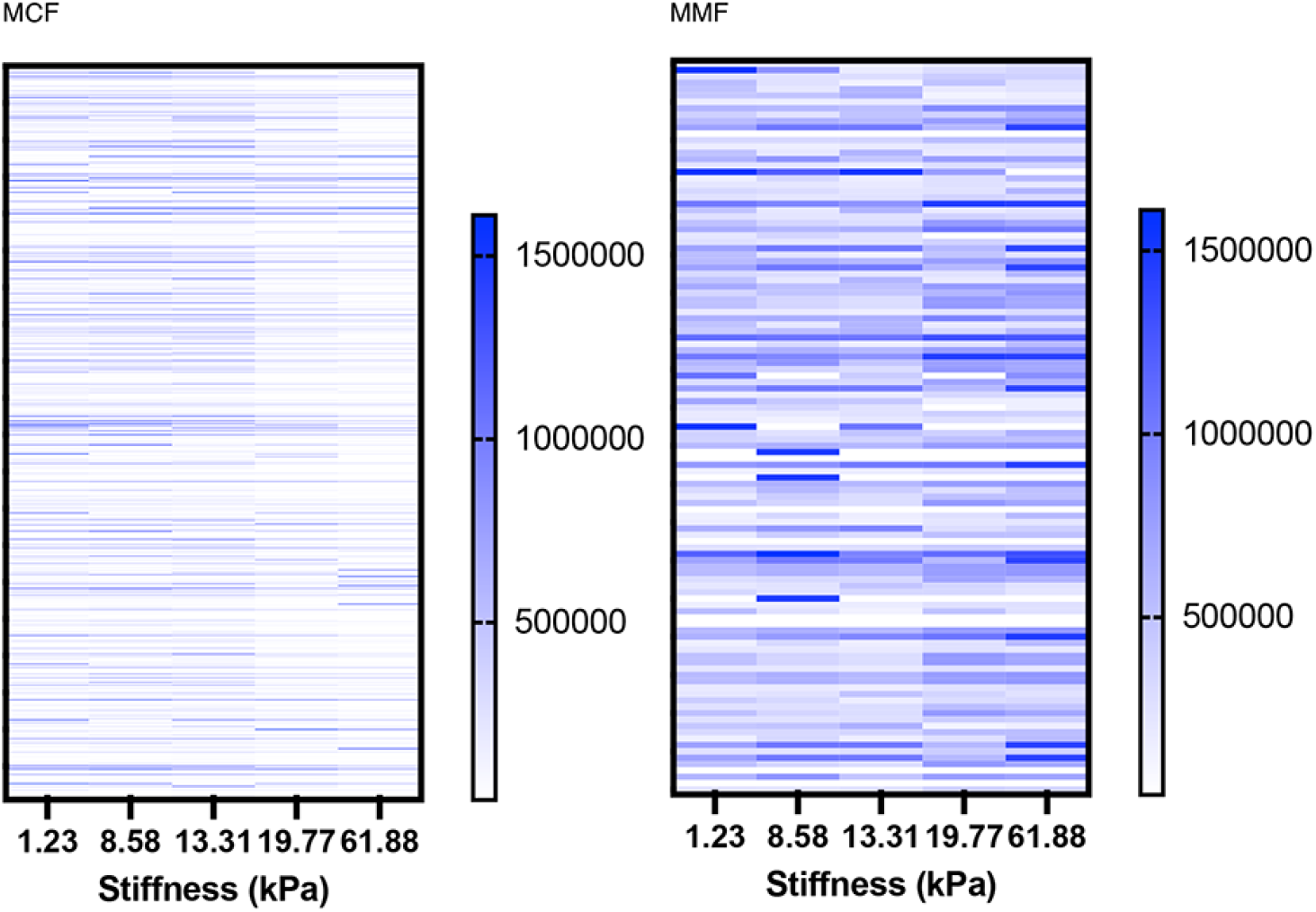
Proteome profile heatmap of stiffness-tuned MMF and MCF secreted EVs.

**Supplementary Figure S4.**
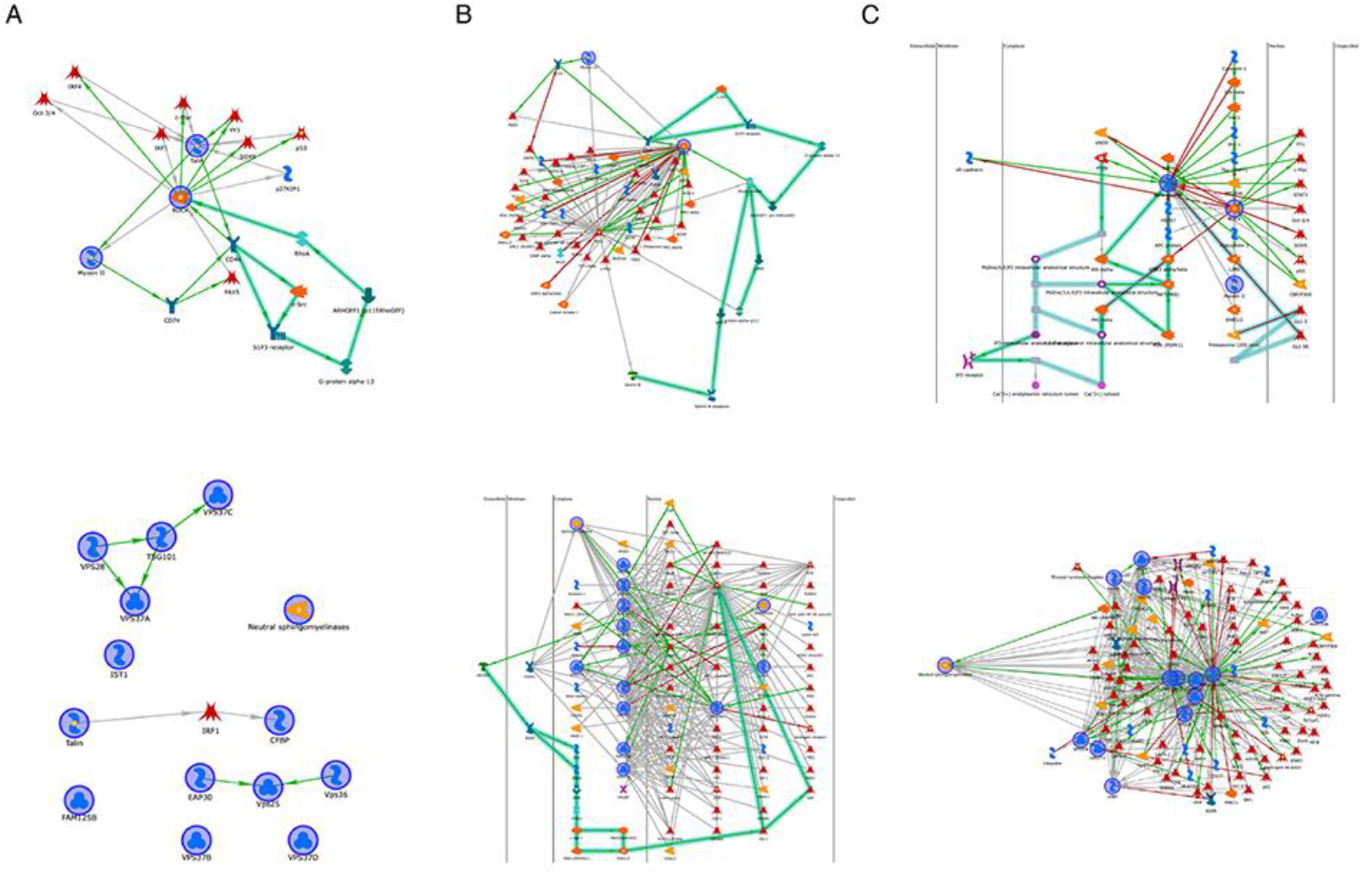
Pathway analysis of identified proteins with EV biogenesis and mechanotransduction pathways. A) Pathway analysis of talin with EV biogenesis and mechanotransduction pathways. B) Pathway analysis of JunB with EV biogenesis and mechanotransduction pathways. C) Pathway analysis of beta-catenin with EV biogenesis and mechanotransduction pathways.

**Supplementary Figure S5.**
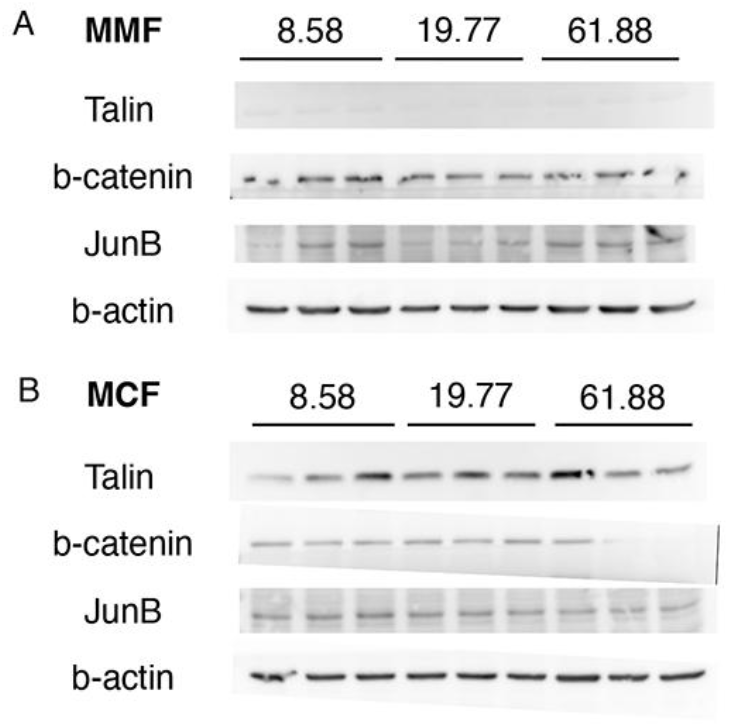
Western Blot analysis of additional proteins in the study. A) Western blot of selected protein expression in MMFs. E) Western blot of selected protein expression in MCFs.

## Notes

Funding Statement: Research reported in this publication was supported by NIH award number 1R01CA275423-01A1.

### Competing Interest Statement

The authors have declared no competing interest.

